# *De novo* transcriptome profiling of mustard aphid (*Lipaphis erysimi*) and differential expression of transcripts associated with developmental stages, feeding and non-feeding conditions

**DOI:** 10.1101/717470

**Authors:** Rubina Chongtham, Kirti Kulkarni, Rohit Nandan Shukla, Gopal Joshi, Amar Kumar, Shailendra Goel, Manu Agarwal, Arun Jagannath

## Abstract

*Lipaphis erysimi* is a Brassicaceae specialist aphid, which causes significant losses in yield and/or reduction of oil content of vegetable and oilseed brassicas and is a major pest in the Indian subcontinent. This study reports the *de novo* transcriptome of *L. erysimi* for the first time. We also present a comparative analysis of nymphs and adult transcriptomes to study the differential expression profiles associated with different developmental stages as well as different feeding conditions. For this, RNA-seq was performed on three different biological samples adults, nymphs (with all nymph stages pooled) and adults starved for 3 hours (referred to as Adult Feeding, AF; Nymphs Feeding, NF, and Adult Starved_3 hr, ANF samples henceforth). A final transcriptome comprising 52,652 transcripts of 1064bp average length and N50 value of 1806 bp was generated. A total of 27,112 transcripts were annotated with insect proteins from SwissProt, of which 4128 transcripts were components of 165 KEGG pathways. A total of 17,296 transcripts were classified based on their Gene Ontology. Potential transcripts for host selection, detoxification, salivary proteins and effectors, molecular chaperones and developmental genes were identified. A total of 23,532 transcripts that remained unannotated were subjected to BLAST against aphid sequences available at AphidBase and a total of 3091 transcripts had hits with sequences of other aphids in the database, out of which 1380 had protein hits. A total of 20441 found to share no homology to any sequence available in the public domain and could therefore represent novel aphid genes or sequences that are unique to *L. erysimi*. This is an exploratory study with no biological replicates. However, the significant repertoire of feeding- and development-related genes and their differential expression profiles generated in this study adds to the limited data available on *L. erysimi* and it would facilitate studies on the molecular basis of aphid feeding and development. This could also allow identification of novel target genes for development of RNAi-based aphid control methods.

## Introduction

Aphids are phloem-feeders of the Order Hemiptera and are one of the most devastating pests of cultivated crops globally. There are over 5000 species of aphids (1) of which, ∼450 species are pests of crop plants (2). The adaptations of aphids to a phloem diet, their ability to overcome plant defences and their rapid rate of multiplication by parthenogenesis, contribute to their successful infestation of crop plants. *Lipaphis erysimi*, also known as mustard aphid, is a Brassicaceae specialist and is one of the most significant challenges to cultivation of the oilseed crop, *Brassica juncea* (Indian mustard) in the Indian subcontinent (3). *B. juncea* accounts for ∼90% of the acreage under rapeseed-mustard cultivation in India (4) and is the second most important oilseed crop worldwide according to the OECD-FAO Agricultural Outlook, 2015-2024. *L. erysimi* not only causes extensive losses in yield (65-96%) and reduction in oil content (by upto 15%), (5, 6) but also reduces crop productivity indirectly by transmitting disease-causing plant viruses (7).

Aphid-plant interactions are complex and initiate with identification and recognition of a host plant based on visual, olfactory and gustatory cues followed by host acceptance and sustained phloem feeding (8-11). In plants, the first line of defence against aphid infestation includes physical barriers viz., a waxy cuticle (12) or glandular trichomes (13). Infested plants may alter expression of genes involved in cell wall metabolism and remodelling (14). When the cell wall is breached, secondary metabolites, such as glucosinolates (15) and enzymes such as proteases (16) and protease inhibitors (17) are released and act upon the invading aphids. Injury to phloem is mitigated by phloem proteins (18) and callose plugs (19). Additionally, plants may release volatiles to attract natural predators of aphids (20) and produce Reactive Oxygen Species (ROS), leading to hypersensitive response (HR). However, aphids have evolved well-adapted mechanisms to counter these defences. They have specialized piercing-sucking mouthparts with long and slender stylets that achieve intercellular penetration and feeding from phloem with minimal physical injury to the plant (21). The aphid saliva lubricates stylet entry through layers of plant tissue (22). Although it induces local as well as systemic plant resistance by releasing elicitors, it can also suppress local plant defence by binding Ca^2+^ ions required for phloem occlusion and oligogalacturonides-induced plant defence (23, 24). Aphid salivary proteins are involved in degradation of plant proteins, detoxification of plant defence molecules and interference with plant signalling pathways (25). Additionally, the ability to compartmentalize glucosinolateshas also been demonstrated in the *Brassica* specialists, *L. erysimi* and *Brevicoryne brassicae* (26).

Expression analyses of genes involved in the above processes can provide important cues to control aphid infestation. Aphids of different developmental stages have been demonstrated to show different transcriptomic responses to host plant in terms of detoxification, apart from expected differences in cuticle formation and hormone-related transcripts (27). Additionally, aphids have been reported to demonstrate high phenotypic plasticity and can transform from wingless to winged forms under conditions of over-crowding, lack of nutrition and upon interactions with other organisms (28, 29). Transcriptome studies have been used to understand the molecular basis of development, aphid-plant interactions and aphid-endosymbiont interactions on the model pea aphid, *Acyrthosiphon pisum*. It has been used to study differential expression of wing development genes (30), role of microRNAs in phenotypic plasticity (31), identification of secretory and effector proteins (25), molecular variations between sexual and asexual morphs (32) and molecular interactions between the aphid and its endosymbiont (33). Trancriptome studies have also been performed in some non-model aphids viz., the soybean aphid (*Aphis glycines*) (34) and grain aphid (*Sitobion avenae*) (35). Changes in expression levels of proteins viz., Heat shock proteins (HSPs) and Cyt P450s was shown to aid adaptation of cotton aphid to different stages of growth and development of its host (36). However, transcriptome studies on plant-aphid interactions have primarily focussed on changes in the host plant transcriptome during compatible and non-compatible interactions with aphids (37-39). A comprehensive analysis of transcriptomic changes within the aphid during such interactions with its host is lacking. Moreover, differential expression of aphid genes during feeding and non-feeding conditions has not been studied until now. There is also limited information on gene expression changes during development from nymphs to adults.

In the present study, we performed *de novo* transcriptome sequencing of *L. erysimi* to investigate differential expression profiles between normally feeding and adults starved for 3h and during different developmental stages of the aphid. To the best of our knowledge, this is the first report on global transcriptomic differences between normal feeding and 3h starvation conditions in any aphid and outlines the possible roles of different sets of genes during feeding conditions (involving host identification, stylet entry and detoxification) and during short non-availability of food. This is also the first study to demonstrate transcriptomic difference between adults and nymphs of *L. erysimi*.

## Materials and methods

### Rearing of *L. erysimi*

*L. erysimi* cultures consisting of parthenogenetic apterous female were initiated from a single cohort of apterous female adults (provided by Prof. A. K. Singh, Department of Zoology, University of Delhi) and propagated in a culture room at 21±1°C with a photoperiodic cycle of 16 hours light: 8 hours dark and a relative humidity (R.H.) of ∼55-70%. The cultures were maintained on excised cauliflower leaves placed over moistened blotting paper in transparent plastic boxes. The leaves were changed every alternate day, distributing progenies to new boxes to prevent nutritional stress and crowding in order to maintain a uniformly apterous culture.

### Collection of biological material

As shown in the schematic representation of the experimental design (Fig 1), three types of biological samples were collected for transcriptome sequencing: (i) Adult Feeding (AF)-comprising adult apterous aphids that were allowed to feed normally (ii) Adult starved for 3h (ANF) and (iii) Nymph Feeding (NF) - comprising normally feeding nymphs of all four nymphal stages (N1 – N4) pooled together. All samples consisted of ∼300-400 apterous morphs. Segregation of the adult and nymphal stages was done under a Stemi DV4 microscope (Zeiss). The AF and NF samples were harvested directly from *B. oleracea* leaves and only those aphids were harvested that had their stylets inserted and were feeding. For the ANF sample, apterous adult aphids were segregated in empty plastic boxes subjecting them to non-feeding conditions for 3 hours. All samples were flash frozen in liquid Nitrogen and stored at −80 °C till further use.

**Fig 1.**
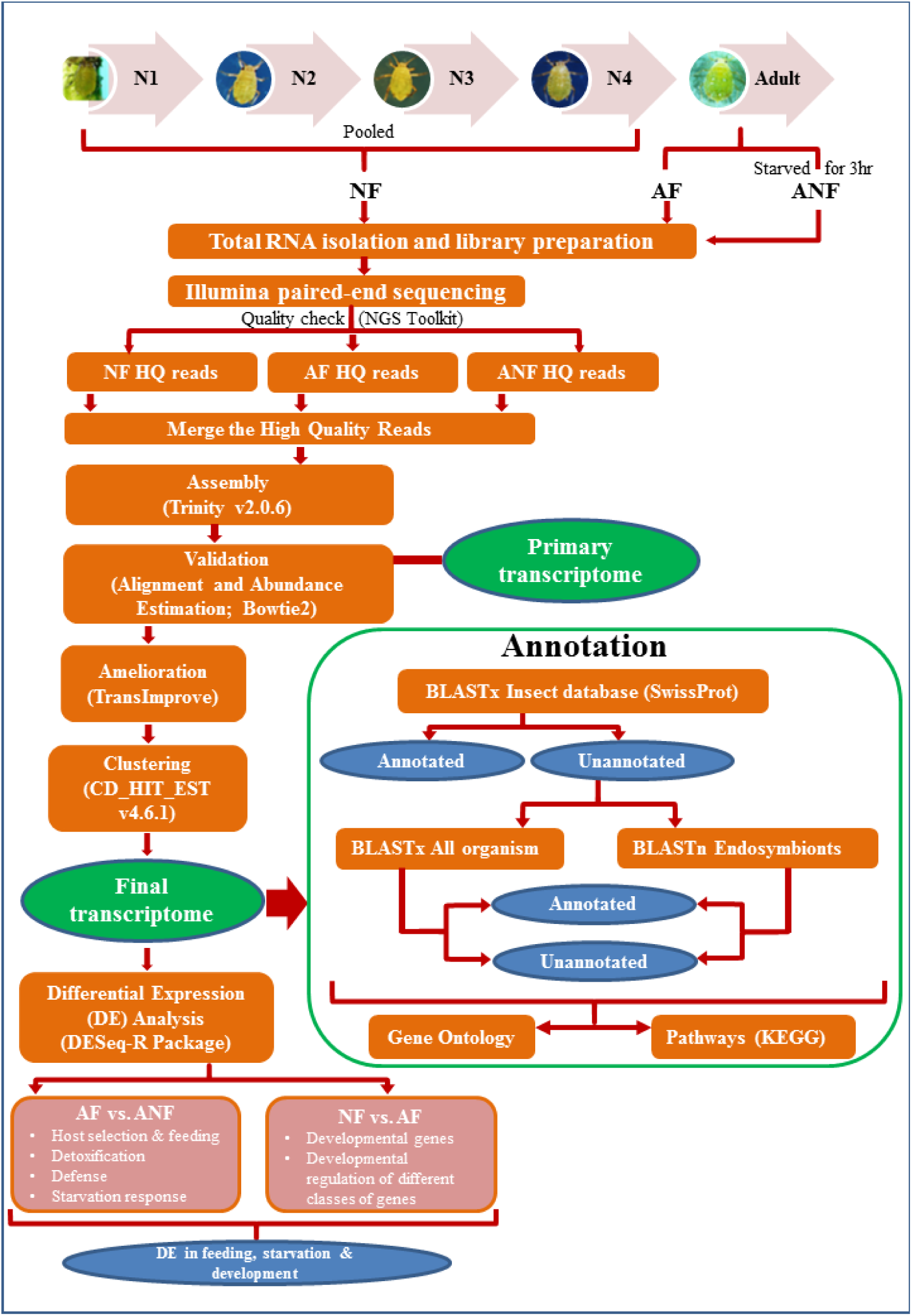
Flowchart representing the complete work-flow for the transcriptome analysis of *L. erysimi*. The tools used at different steps are given within parentheses. N1-4: nymphal stages 1 to 4, NF: Nymph Feeding, AF: Adult Feeding, ANF: Adult starved for 3h and HQ: High Quality.

### Preparation of cDNA libraries, sequencing and data analysis

Total RNA was isolated from each sample using Trizol method (40). cDNA libraries were prepared from 2µg total RNA per library using TruSeq RNA Sample Prep Kit v2 (Illumina Inc., USA) according to manufacturer’s instructions. The quality of cDNA libraries was assessed using 2100 Bioanalyzer (Agilent Technologies). The libraries were sequenced with Genome Analyzer IIx and HiSeq 2000 (Illumina, USA) sequencer using paired-end method with an expected average read length of ∼100bp. The raw reads were subjected to quality check using NGS QC toolkit (41), in which, adapter sequences were trimmed and low quality reads (with read length <70% and a phred score <20) were removed. The resulting high quality filtered paired sequence reads from Adult Feeding, Adult Non-Feeding and Nymph Feeding samples were pooled and provided as an input for transcriptome assembly. *De novo* transcriptome assembly and validation was performed using Trinity assembler (42). The primary transcriptome assembly thus obtained was validated by mapping the HQ filtered reads back to the assembled reads using Bowtie2 tool (43). Transcript abundance was estimated for each sample using RSEM tool (44). The transcriptome data was further ameliorated using TransImprove (Bionivid Technology Pvt. Ltd., Bengaluru), which analysed transcriptome integrity on the basis of transcript depth and coverage that were calculated based on read length and total transcript length (45) to generate the final transcriptome assembly. The final transcriptome was clustered with CD-HIT-EST (v4.6.1) (46). Normalization of the assembled transcriptome, differential gene expression analysis and biological significance analysis was done using DESeq-R package (47). Fold-change of transcripts for AF vs. ANF and NF vs. AF were calculated and converted to log_2_ scale. Transcripts with log_2_ fold-change ≥ +2 and ≤ −2 with p ≤ 0.05 were considered as significantly up/down-regulated. Annotation for final transcripts longer than 200 bp was done using BLASTx homology search (48) against all Insect Protein sequences from SwissProt. The remaining unannotated transcripts were aligned against protein sequences of all organisms in the NCBI non-redundant database using BLASTx. BLASTn was performed for the unannotated transcripts against sequences of bacterial endosymbionts (listed in *Buchnera*Base) in the NCBI nucleotide database. Blast hits were filtered for transcripts with E-value scores ≤ 1e-3. Gene Ontology (GO) annotation was done and categorized with respect to Biological Process, Molecular Function, and Cellular Component. Pathway analysis of the transcripts was performed based on the Kyoto Encyclopedia of Genes and Genomes (KEGG) database. The transcriptome was analyzed to identify transcripts with one, two or all of these: signal peptides (SignalP4.1), transmembrane domains (TMpred) and nuclear localization signal (SeqNLS).

### Real Time quantitative PCR

Total RNA was isolated from each of the three samples described above in three independent biological replicates. Five µg of each RNA sample was treated with 2U DNAseI (New England Biolabs) at 37°C for 30 minutes followed by Phenol-Chloroform-Isoamyl alcohol extraction, Sodium acetate-Ethanol precipitation and wash with 70% Ethanol. RNA integrity was checked on a 1.2% denaturing agarose gel. cDNA was prepared from 500 ng of each DNA-free RNA sample using iScript Reverse transcription kit (Invitrogen) following manufacturer’s protocol. After testing several candidate genes for use as an endogenous control, the pea aphid *Elongation Factor 1α* (*EF1α*) was found to show stable expression levels in all the three samples of the current study and was used for normalization. The *EF1α* primers used were: 5’ ATGGAATGGAGACAACATGTTGG 3’ (forward primer) and 5’ AGAGCCTTGTCAGTTGGGCG 3’ (reverse primer). PCRs were done on a CFX Connect Real-Time System (BioRad) in 96-well plates using SYBR Green (Roche) for selected transcript sequences with two technical replicates for each of the three independent biological replicates. Primer sequences for validated transcripts are provided in S1 Table. ΔΔCt was calculated to determine the relative expression of each transcript (49).

## Results and discussion

### Analyses of sequencing data and *de novo* transcriptome assembly

Three different biological samples of apterous aphids viz., AF, ANF and NF were collected and used for construction of transcriptome sequencing libraries. While the AF and NF samples were collected directly from *B. oleracea* leaves (as described in the Materials and Methods), the ANF samples were starved for 3 hours prior to harvesting. It has been shown in earlier studies that the threshold duration of phloem feeding indicating acceptance of the host is ≥10 minutes (50). An earlier study on *M. persicae*, which is a generalist aphid that also infects Brassicas, showed that effects of nutrition loss on aphid growth due to starvation for <4 hours was overcome once returned to host plants but not when the starvation period exceeded four hours (51). Thus, an induced non-feeding state by starvation for 3 hours was followed in this study to ensure arrest of aphid activities involved in aphid-host interactions without causing irreversible damage to aphid health and survival.

As given in Fig 1, the three transcriptome libraries were sequenced using high-throughput Illumina sequencing platform and resulted in 622.87 million raw paired end reads (Table 1). Low quality reads with >70% of the bases having a phred score <Q20 were filtered and removed. Further, adapter sequences were trimmed resulting in 536.40 million high quality (HQ) filtered reads, spanning a total length of 53.96 billion bases. The number (in millions) of HQ filtered reads obtained were 165.69, 155.29 and 215.41 in AF, ANF and NF samples, respectively (Table 1).

**Table 1.**
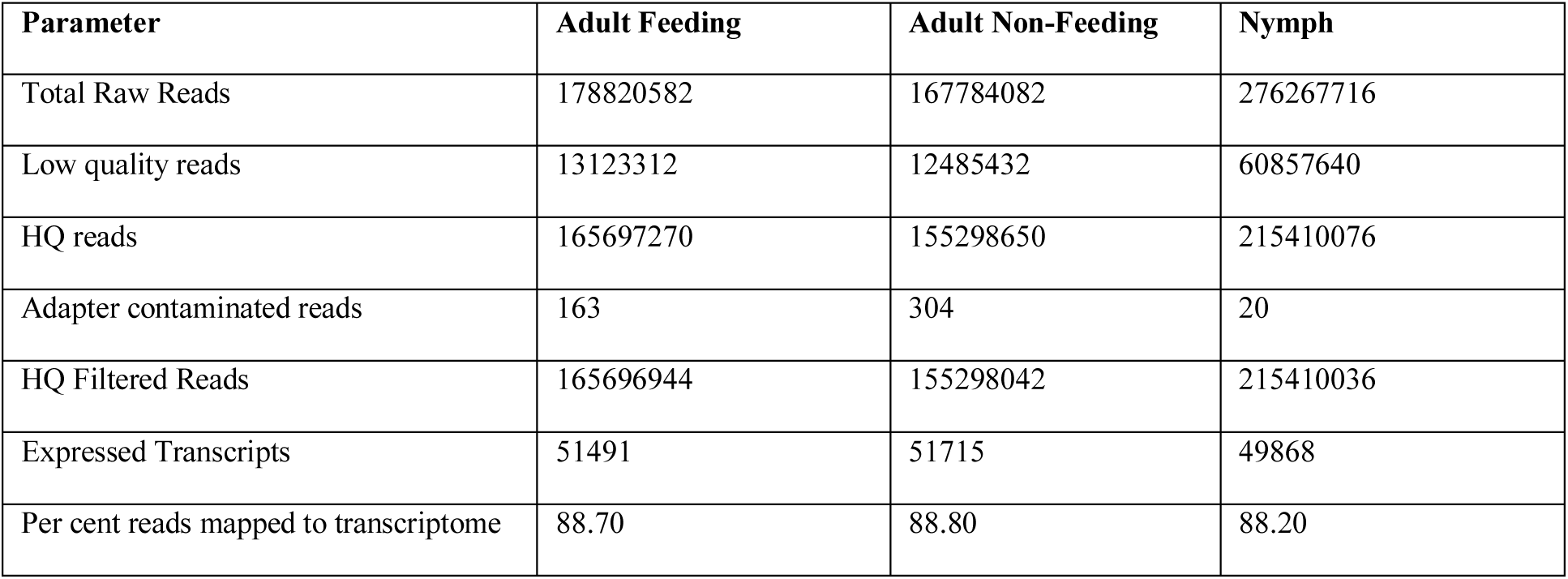
Sequencing data and quality analysis of the L. erysimi transcriptome.

Due to lack of genome sequence information of *L. erysimi*, a *de novo* assembly of the transcriptome data was performed using Trinity assembler (42). The HQ reads from all the samples (536.40 million) were merged to generate the transcriptome assembly. The primary assembly was subjected to further correction by mapping HQ reads back onto the assembly, resulting in ∼88% reads getting mapped. The resulting primary transcriptome yielded a total of 110,556 transcripts spanning a length of 89 million bp with a transcript length ranging from 224bp to 26824bp. The average transcript length was 807 bp with an N50 value of 1475 bp. Transcriptome integrity was analysed using TransImprove 2.0.1 (Bionivid Technologies, India) and transcripts with a coverage of ≥ 70% and a minimum depth of 5X were selected for further analysis. This step was performed to increase the sensitivity and specificity of transcriptome assembly (45). The contigs from each sample were clustered and redundancy was removed, resulting in a final ameliorated transcriptome of 56 million bp with 52,652 transcripts, 31.78% of which were ≥ 1000bp in length. A comparison of the primary transcriptome and the final ameliorated transcriptome is given in Table 2. The amelioration step improved the average transcript length and the N50 value of the final transcriptome to 1064 bp and 1806 bp, respectively. The average G+C content of the final *L. erysimi* transcriptome was low at 34.52%, which is in consonance with earlier studies on insect genomes (52, 53). All the sequencing data has been submitted to the Sequence Read Archive of NCBI with accession number SRP093554.

**Table 2.** Assembly statistics of high quality reads.

| Parameter | Primary Assembly Values | Final Assembly Values |
| --- | --- | --- |
| Number of Final Transcripts | 110556 | 52652 |
| Final Transcriptome Length (bp) | 89232845 (~89Mbp) | 56027788 (~56 Mbp) |
| Minimum Transcript Length(bp) | 224 | 224 |
| Maximum Transcript Length(bp) | 26824 | 26824 |
| Average Transcript Length(bp) | 807.13 | 1064.12 |
| N50 | 1475 | 1806 |
| (G+C)% | 34.75 | 34.52 |

### Annotation of assembled transcripts of *L. erysimi* and its endosymbiont

The 52,652 transcripts were subjected to BLASTx-based annotation against insect protein sequences in Swiss-Prot database. A total of 27,112 transcripts (50.5%) were annotated with insect proteins from SwissProt (S1 Data sheet), of whcih 96.13% had E-values ≤ 10^−5^. More than 86% of these insect-annotated transcripts (23,429) comprised of *A. pisum* sequences. The distribution of the annotated transcripts among the top ten organisms is given in S2 Table. The remaining 25,540 unannotated transcripts were subjected to two independent BLAST analyses: BLASTx against NCBI non-redundant (nr) database comprising all organisms and BLASTn against nucleotide sequences of sequenced endosymbionts listed in *Buchnera*Base. This resulted in the annotation of 816, 128 and 1064 transcripts with endosymbionts (including 336 sequences from *Buchnera aphidicola*; S2 Data sheet), *A. pisum* and sequences of other organisms (S3 Data sheet), respectively. A total of 23,532 transcripts did not show homology with any sequence available in public domain and may therefore represent novel aphid transcripts and/or sequences that are unique to *L. erysimi*.

The 52,652 assembled transcripts were subjected to Gene Ontology analysis resulting in classification of 17,296 transcripts by at least one of the three ontologies i.e., biological process, molecular function and cellular component. Pathway annotation was done using KEGG Database and 4128 insect-annotated transcripts could be assigned to 165 different KEGG pathways. A significantly higher number of transcripts (1123, 27.2%) belonged to the category of metabolic pathways than others such as ribosome (257, 6.2%), RNA transport (190, 4.6%), spliceosome (138, 3.3%) and protein processing in endoplasmic reticulum (136, 3.3%). The top 10 pathways are given in Table 3.

**Table 3.** Top five enriched KEGG pathways in order of abundance.

| KEGG ID | Pathway | No. of transcripts |
| --- | --- | --- |
| ko01100 | Metabolic pathways | 1123 |
| ko03010 | Ribosome | 257 |
| ko03013 | RNA transport | 190 |
| ko03040 | Spliceosome | 138 |
| ko04141 | Protein processing in endoplasmic reticulum | 136 |

### Identification and differential expression of transcripts based on feeding status (feeding vs. non-feeding adults) and developmental stages (nymphs vs. adults)

Pairwise differential expression analysis of whole transcriptomes of AF, ANF and NF samples identified genes that are differentially expressed in different developmental stages (NF vs. AF) and under different feeding conditions (AF vs. ANF). The distribution of transcripts among the three biological samples is shown in Fig 2. For identification of differentially expressed transcripts in the studied samples, transcripts with Log_2_ Fold change ≥ 2 or ≤ −2 were considered as significantly up-regulated or down-regulated, respectively between the samples.

**Fig 2.**
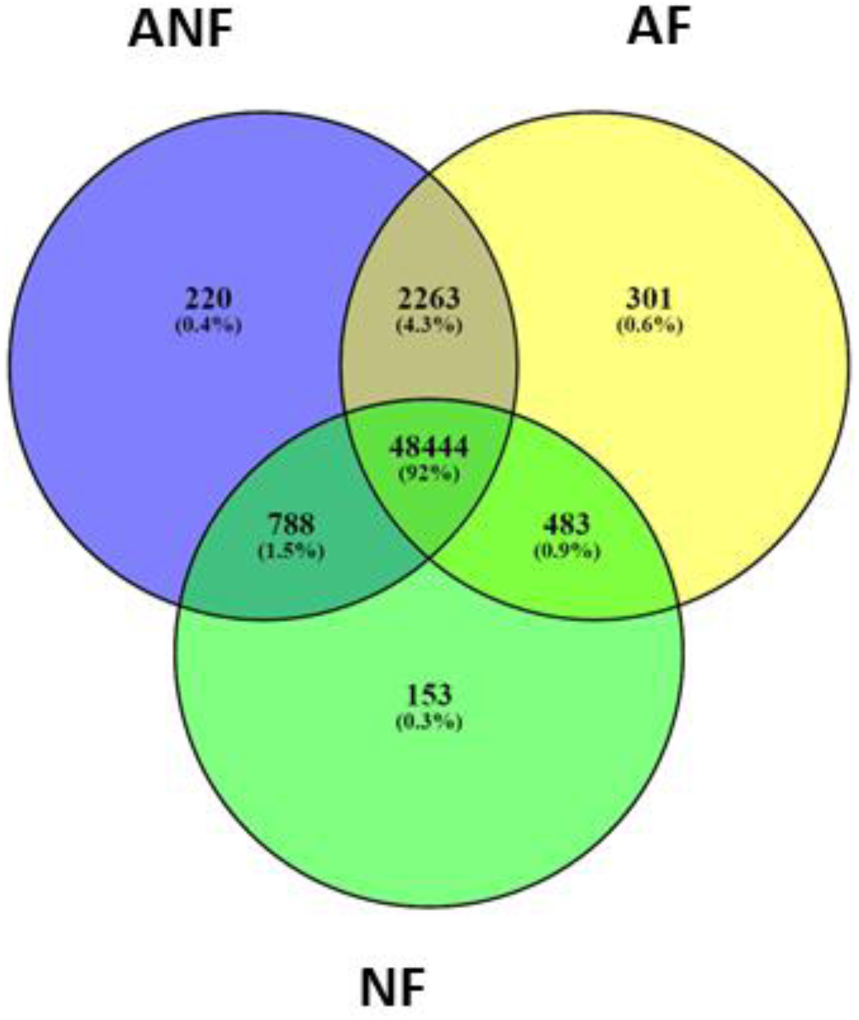
Venn diagram showing distribution of transcripts in Adult Feeding (AF), Adult starved for 3h (ANF) and Nymph Feeding (NF) samples.

The distribution of up-regulated and down-regulated transcripts is shown in Figs 3a and 3b. An unsupervised hierarchical clustering of differentially expressed transcripts using Pearson Uncentered algorithm with average linkage rule showed distinct patterns of up-regulated and down-regulated genes among the three samples (Figs 3c, 3d). These results are described and discussed in detail in the following sub-sections.

**Fig3.**
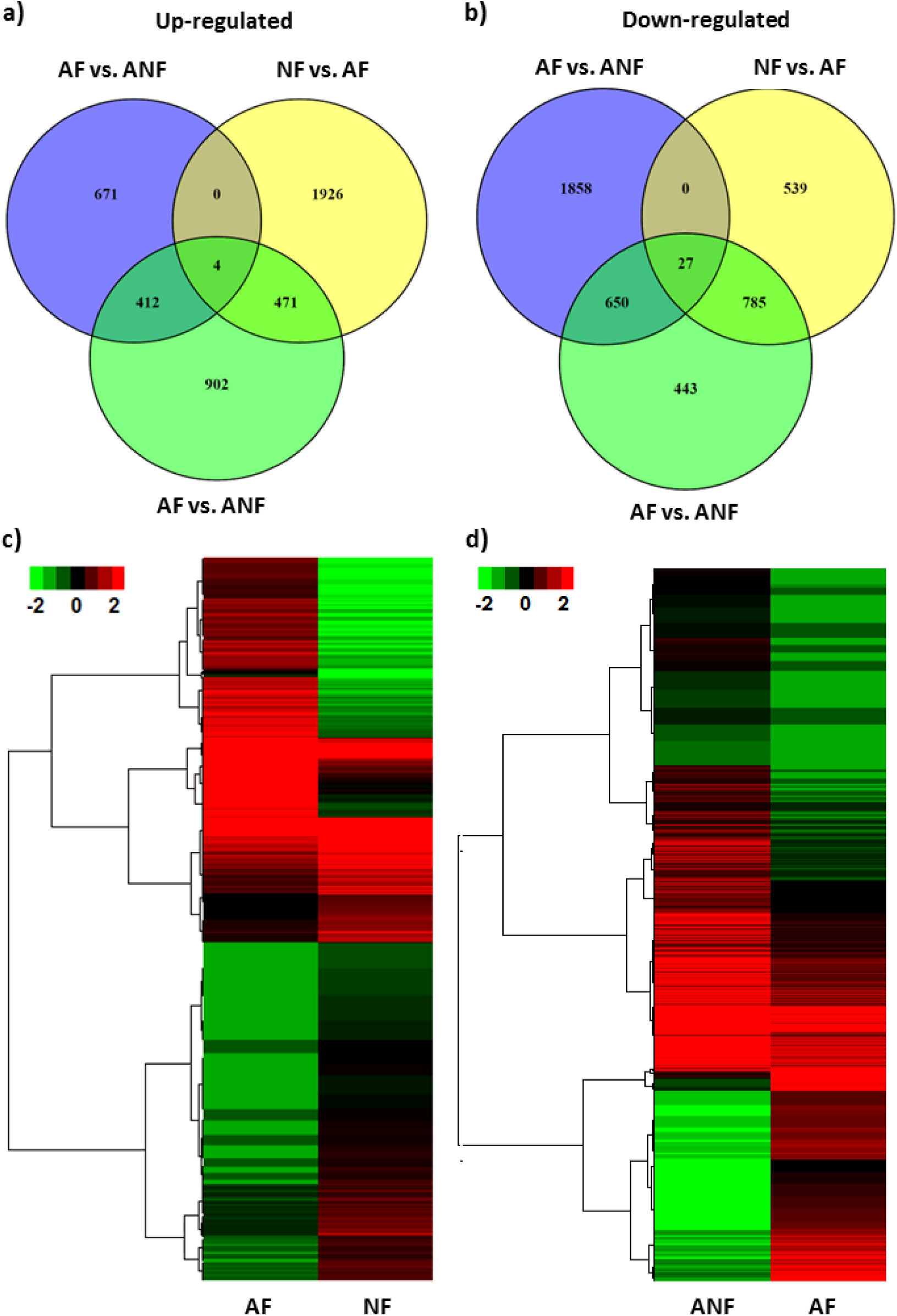
Expression analysis of differentially expressed transcripts in *L. erysimi* transcriptomes. Venn diagrams representing the number and distribution of transcripts in the comparisons between (**A**) AF vs. ANF and (**B**) NF vs. AF. Heat maps with colour scales representing an unsupervised hierarchical clustering of differentially expressed transcripts between (**C**) AF & NF and (**D**) ANF & AF. AF: Adult Feeding, ANF: Adult starved for 3h, NF: Nymph Feeding.

### Differentially expressed transcripts under varying feeding conditions

In the comparison between AF and ANF samples, 1087 transcripts showed higher levels of expression in feeding adults (Fig 3a; S4 Data sheet) and 2535 transcripts showed higher expression in non-feeding adults (down-regulated in AF vs. ANF; Fig 3b; S5 Data sheet). There were 784 and 1008 transcripts that were ‘specific’ to feeding and non-feeding aphids, respectively (S6 and S7 Data sheets, respectively). It may be noted here that biologically, the ‘specific’ transcripts might be highly up-regulated (or down-regulated) transcripts in the relevant categories.

#### Biological processes influenced by feeding status and differential expression of associated genes

The major biological processes influenced by feeding status are given by the ten most abundant GO terms for Biological Processes among the up-regulated transcripts in feeding and non-feeding aphids and among the specific transcripts (Figs 4a and 4b, respectively).

**Fig 4.**
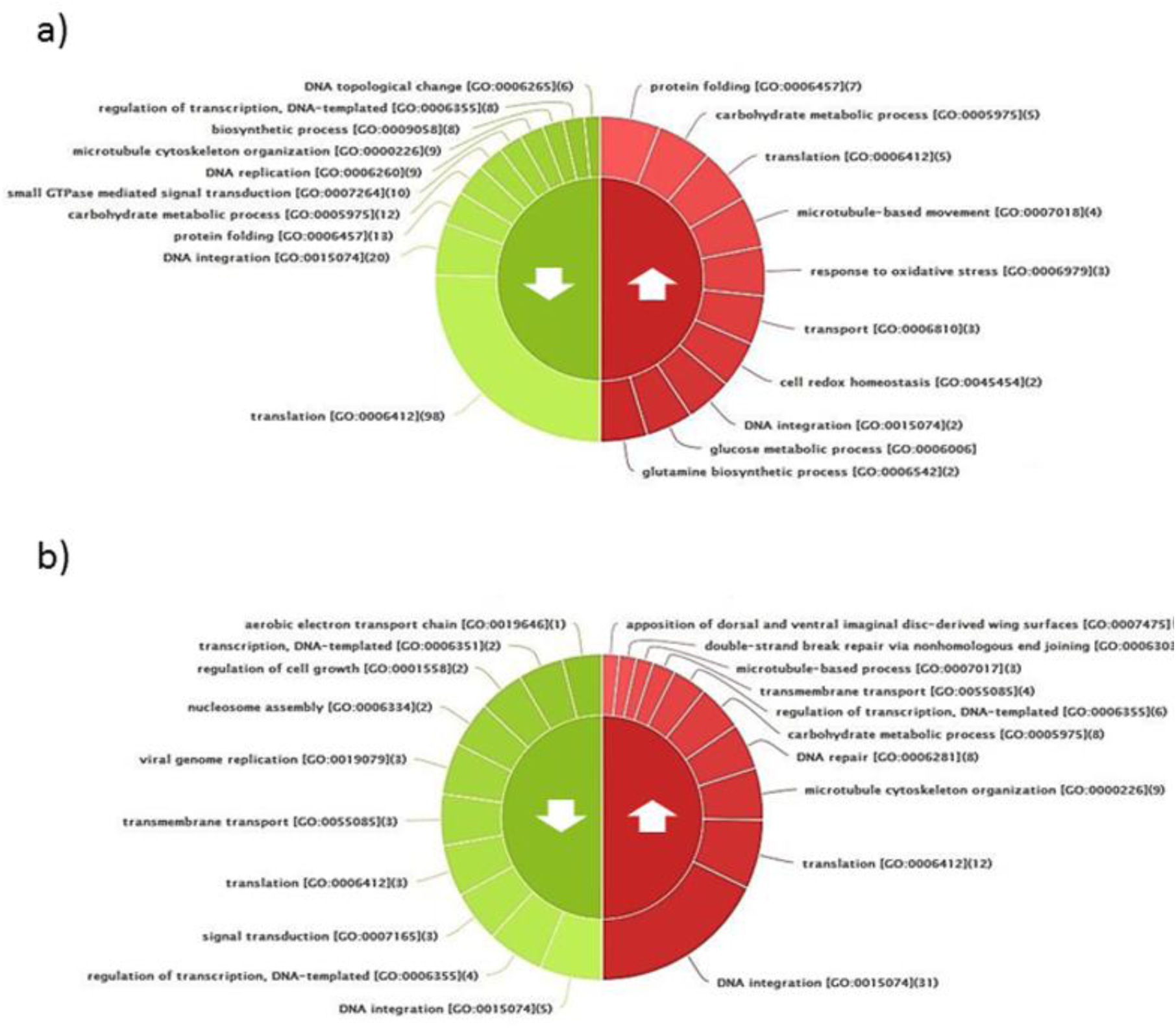
Classification of enriched differentially expressed transcripts on the basis of Gene Ontology: Biological Process. Top 10 enriched transcripts are represented for the comparison between (**a**) AF & ANF, and (**b**) NF & AF.AF: Adult Feeding, ANF: Adult Non-Feeding, NF: Nymph Feeding. Up and down arrows in the figure represent the up-regulated and down-regulated transcripts respectively.

“Protein folding” was one of the ten most abundant biological processes identified among the up-regulated transcripts in both feeding and non-feeding aphids and among those specific to feeding aphids in comparison to non-feeding aphids. All transcripts involved in protein folding were either HSPs or other chaperones. Out of the seven up-regulated transcripts involved in protein folding, two were of HSP 83, an HSP 90 homolog, which plays pleiotropic roles in longevity, fecundity and embryogenesis of *A. pisum* (54). On the basis of BLASTx analysis, a total of 35 Hsps were identified in our study, including four that were specific to feeding adults in addition to seven up-regulated ones. In non-feeding adults, thirteen transcripts involved in protein folding were up-regulated of which, ten were known chaperones. Subjecting the aphids to starvation for 3 hours to induce a non-feeding state could have induced stress leading to higher expression of chaperones probably to cope with starvation stress. The roles of bacterial HSPs in the interactions of aphids with their biotic/abiotic environment have been described in earlier studies (55) and our study reveals the possibility of aphid chaperones playing an important role during feeding and starvation as well. Though the GO term, ‘protein folding’ is abundant among the up-regulated transcripts in both feeding and non-feeding aphids, it is interesting to note that different sets of chaperones (except calreticulin) are up-regulated in each sample. This suggests a differential role of molecular chaperones during feeding and non-feeding conditions. Transcripts of molecular chaperones with their expression patterns are given in S3 Table.

“Carbohydrate metabolism” is one of the most abundant biological processes among the feeding-specific, non-feeding-specific and up-regulated as well as down-regulated transcripts. Out of the five up-regulated transcripts in feeding aphids, three were putative Glycosyl hydrolase family 18 (GH18), two of which, had signal peptide sequences. GH18 has earlier been identified as a potential effector in plant-fungus interaction (36). The other two transcripts were glycogen phosphorylase and sucrase. Transcripts involved in carbohydrate metabolism and specific to feeding aphids were associated with enzymes viz., Fructose-1, 6-bisphosphatase, Polygalacturonase and Chitinase. With the availability of food, higher expression of glycolytic enzyme such as Fructose-1, 6-biphosphate is expected. Polygalacturonase on the other hand, is a cell wall degrading pectinase and presumably plays a role in disrupting the plant cell wall in order to facilitate penetration. An earlier study had reported the presence of polygalacturonase in the saliva of *Schizaphis graminum* (56) and our results indicate direct involvement of this enzyme during feeding.

Another top GO term among up-regulated transcripts of feeding aphids was “Response to oxidative stress”. Earlier studies have shown enhancement of detoxification enzymes in aphids in response to artificially elevated stressors in plants (57, 58). In our study, we found that such enzymes are over-expressed during routine feeding on host plants as well. Three such transcripts encoding catalase and cytochrome c peroxidise were identified. Catalase and cytochrome c peroxidise are involved in managing reactive oxygen species (ROS) as components of antioxidant defence mechanism (59, 60). Their enhanced expression might play an important role in preventing hypersensitive response of the plant to counter aphid feeding or oxidative stress from the ingested plant material. Other classes of proteins associated with response to oxidative stress are Glutathione-S-transferases (GSTs) and Superoxide dismutases (SODs). In our study, three GST transcripts were detected of which, one transcript showed higher expression in non-feeding aphids. Of the eleven SOD transcripts identified in this study, three showed up-regulated expression in non-feeding aphids.

The GO biological process, “Transport” also showed positive differential regulation under feeding conditions. Up-regulated transcripts were twelve in number and were involved in transport (3 transcripts), transmembrane transport (2 transcripts), vacuolar transport (2 transcripts) and other transport (5transcripts). Among the feeding-specific transport-related transcripts, two were involved in transmembrane transport and three more in other types of transport. Twenty transcripts with GO term “DNA integration” (GO: 0015074) that are associated with transposition/ DNA recombination were up-regulated in non-feeding aphids, in contrast to only two transcripts that were up-regulated in feeding aphids (Figs 4a & b). This might be an indication that transposition or DNA recombination are probably inducible by a short period of starvation stress. The implications of this observation, which needs further investigation for confirmation, have been discussed in later sections.

Among the transcripts specific to non-feeding adults, other enriched metabolic processes include gluconeogenesis, glucose and glycogen metabolic processes and glycolytic process. This indicates that storage glycogen is mobilized for nutrition in the non-feeding aphids. Interestingly, the enriched chitinase activity along with other energy metabolic processes is accompanied by the up-regulation of developmental genes, viz., Homeobox proteins genes (*goosecoid*, *PKNOX2* and *Prospero*), *Mothers against decapentaplegic* homolog (MAD homolog) and *Down Syndrome Cell Adhesion Molecule* (*DSCAM*) splice variant 3.12.3.1, which were specific to non-feeding aphids (S7 Data sheet). MAD homolog mediates the function of *Decapentaplegic* (61), which is a wing developmental gene of *A. pisum* (30). The activity of the developmental genes may be explained as a probable effect of starving ∼300 aphids together for 3 hours during sampling that may introduce starvation and crowding effect. Both starvation and crowding are known to induce wing development in aphids (28, 62, 63).

#### Differential expression of genes in other feeding-related functional categories

In order to identify other transcripts that are required for aphid feeding on host plants, we analyzed the functional categories of proteins involved in host recognition and detoxification. Of these, host recognition and detoxification related transcripts that were differentially regulated in feeding and non-feeding aphids are given in **Table 4** (complete list is given in S4Table). Odorant Binding Proteins present in their antennae (64) relay olfactory signals to odorant receptors (ORs), thereby aiding host recognition. Out of 31 ORs identified in our study, three were up-regulated in feeding aphids as compared to non-feeding ones. Only one OR transcript was up-regulated in non-feeding aphids. Another relevant functional category included detoxification-related proteins viz, cytochrome P450 and laccase. We identified 14 cytochrome P450 transcripts and 4 NADPH-cytochrome P450 reductase transcripts in our transcriptome data. Only one NADPH-cytochrome P450 reductase transcript was up-regulated in feeding adults w.r.t. non-feeding adults. Cytochrome P450 6a9 and Cytochrome P450 6B5 were specific to feeding aphids. Cytochrome P450 has been earlier shown to increase nicotine resistance in *M. Persicae* (65). The multi-copper binding enzyme Laccase, present in hemipteran salivary glands, oxidises plant phenolics (66). *Laccase 1* has been shown to be up-regulated in *A. glycines* fed on resistant plants (67). We identified transcripts encoding other laccases viz., laccase 2 (fragment), laccase 4, laccase 5 and laccase 25 in *L. erysymi*. While laccase 25 was specific to feeding adults, laccase 2 and 5 were significantly up-regulated in feeding adults in comparison to non-feeding adults. The functional relevance of these transcripts in detoxification of plant toxins needs to be established following which, they could be selected as potential targets for introduction of aphid resistance in crops.

**Table 4.** Transcripts of host recognition and detoxification differentially regulated in feeding and non-feeding aphids.

| Transcript_ID | log<sub>2</sub>FoldChange<br>(AF vs ANF) | Protein Name |
| --- | --- | --- |
| **Odorant receptor** |  |  |
| TR40368\|c0_g1_i1 | -2.00 | Odorant receptor |
| TR18592\|c0_g1_i2 | 2.32 | Odorant receptor |
| TR35268\|c0_g5_i1 | 2.00 | Odorant receptor |
| TR35268\|c0_g2_i1 | 3.70 | Odorant receptor |
| **Cytochrome P450s** |  |  |
| TR59732\|c0_g1_i1 | -2.57 | NADPH--cytochrome P450 reductase |
| TR59884\|c0_g1_i1 | -3.32 | Cytochrome P450 CYP307A1 |
| TR65275\|c0_g1_i1 | -2.22 | Cytochrome P450 CYP6DJ1v3 |
| TR11431\|c0_g1_i1 | -2.81 | Cytochrome P450 (Fragment) |
| TR34267\|c0_g1_i1 | 4.52 | NADPH--cytochrome P450 reductase (EC 1.6.2.4) |
| TR13148\|c0_g1_i1 | AF specific | Cytochrome P450 6a9 |
| TR27685\|c0_g1_i1 | AF specific | Cytochrome P450 6B5 |
| TR42639\|c0_g1_i1 | ANF specific | Cytochrome P450 4C1 |
| **Glutathione-S transferase** |  |  |
| TR29312\|c0_g1_i1 | -3.85 | Glutathione-S transferase (Fragment) |

| **Superoxide dismutase** |  |  |
| --- | --- | --- |
| TR3266\|c0_g1_i1 | -2.84 | Superoxide dismutase [Cu-Zn] (EC 1.15.1.1) |
| TR66964\|c0_g1_i1 | -2.03 | Superoxide dismutase (EC 1.15.1.1) |
| TR47568\|c0_g1_i1 | -2.12 | Superoxide dismutase [Cu-Zn] (EC 1.15.1.1) |
| **Peroxiredoxin** |  |  |
| TR36976\|c0_g1_i1 | 2.81 | Peroxiredoxin-like protein |
| **Laccases** |  |  |
| TR35389\|c0_g1_i1 | 3.21 | Laccase-5 |
| TR35148\|c1_g1_i1 | 5.25 | Laccase 2 (Fragment) |
| TR809\|c0_g1_i1 | Feed Specific | Laccase-25 |
| **Cytochrome c peroxidase (mitochondrial)** |  |  |
| TR52985\|c0_g1_i1 | -2.44 | Cytochrome c peroxidase, mitochondrial |
| TR35636\|c0_g1_i1 | 4.60 | Cytochrome c peroxidase, mitochondrial |
| TR48754\|c0_g1_i1 | Feed Specific | Cytochrome c peroxidase, mitochondrial |

#### Differential expression of putative effectors

During feeding, aphids produce two kinds of saliva: gelling sheath saliva and liquid saliva. Gelling saliva protects the stylet and closes punctured cellular sites during its penetration inhibiting activation of plant defence responses (11). Liquid saliva contains several proteins that function as effectors and induce plant defence response and also proteins that counter plant defence. To identify transcripts encoding putative effector proteins, we analysed the transcriptome for transcripts containing signal peptides and without transmembrane domain or nuclear localization signal. We identified 901 such transcripts of putative effectors annotated with insect proteins of which, four transcripts showed up-regulated or specific expression in feeding aphids compared to non-feeding aphids. These included two transcripts of GH18 domain containing protein and two mosquito proteins with oxidoreductase activity. Both these classes of enzymes have been reported in aphid saliva and/or effectors in other aphids. The Glycosyl hydrolase family 18 includes insect chitinases, whose orthologs have been reported in *A. pisum* (68). Chitinase with signal peptide has also been identified in the saliva of *Diuraphis noxia* by proteomic analysis and might act as an effector (69). Oxidoreductases (of GMC family) have been identified in saliva of *A. pisum*, *S. avenae* and *Metopolophium dirhodum* and have been implicated in detoxification of plant defence molecules (70). Differential regulation of these putative effector transcripts in different developmental stages is discussed later. The complete list of differentially expressed transcripts of putative effectors with annotation is given in S5 Table.

### Differentially expressed transcripts between Nymphs and Adults

A comparison of the nymph and adult transcriptomes identified 2401 transcripts that showed increased expression in nymphs(Fig 3a; S8 Data sheet)and1351 transcripts with higher expression in adults (down-regulated in NF vs. AF; Fig 3b; S9 Data sheet). A total of 941 transcripts were found specific to nymphs and 2564 transcripts were specific to adults (S10 and S11 Data sheets, respectively). The major biological processes differing across developmental stages are indicated by the ten most abundant GO terms for biological processes for these transcripts (Fig4b). The nymph-specific and nymph up-regulated transcripts were predominantly involved in active metabolism, translation of proteins, DNA repair, cytoskeletal organization and developmental processes. Nymphs are metabolically very active and have actively dividing cells. This explains the up-regulation of transcripts associated with developmental processes, microtubule organization and carbohydrate metabolism, along with those of DNA integration and translation.

DNA integration and translation were found to be most significantly influenced by developmental stage of the aphids in this study, both in the nymph-specific and up-regulated categories. ‘DNA integration’ transcripts might be involved in DNA recombination. Transcripts involved in GO biological processes ‘DNA integration’, ‘DNA repair’ and ‘double-strand break (DSB) repair via non-homologous end joining’ (NHEJ) were up-regulated in nymphs. Up-regulation of processes related to DNA repair in nymphs is expected as they are developmentally highly active. DNA repair by both homologous recombination (HR) and NHEJ is known to be regulated spatially and temporally during neuronal development in mammals, with HR being more active during early stages of development (71). Only five transcripts for DNA integration were found to be up-regulated in adult nymphs.

Interestingly, 841 up-regulated transcripts in nymphs (including transcripts for development and chaperones among others; S8 Data sheet), were also up-regulated in non-feeding adults when compared with Adult Feeding sample. Similarly, out of the 941 nymph-specific transcripts in NF vs. AF (Fig 2; S10 Data sheet), 788 transcripts were also expressed in non-feeding adults. This might indicate the activation of developmental genes due to onset of starvation as discussed earlier. This overlap may also be due to the fact that moulting nymphs frequently stop feeding and hence, may show effects similar to starvation (72). The 1351 genes that were down-regulated in nymphs in comparison to adults (i.e., up-regulated in adults) (S9 Data sheet) were distributed across various biological processes, viz., DNA integration, regulation of transcription, signal transduction and membrane transport among others (Fig 4b). The annotated down-regulated transcripts included those coding for Myrosinase 1, for defence against fungi and predators and possibly involved in alarm-system against predators (26), odorant receptor and chemosensory protein for host recognition and others like cell wall-associated hydrolase, salivary protein and secreted proteins. This suggests that colony defence and finding a suitable host and/or feeding-site might be prominent roles for adult aphids rather than nymphs. Of these 1351 down-regulated transcripts, 1165 transcripts were expressed at basal levels (Log_2_ Fold change <2 and >−2) in Adult Feeding vs. Adult Non-Feeding samples (S9 Data sheet). These transcripts might be required in the adult phase irrespective of feeding or starvation conditions and may potentially be involved in adult morphology and/or behaviour. We identified 2564 adult-specific transcripts (S11 Data sheet) for which, the top 5 enriched GO-biological process terms were translation, carbohydrate metabolism, protein folding, microtubule-based process and cell redox homeostasis.

#### Developmental regulation of transcripts related to host identification, detoxification and chaperone activity

Some transcripts belonging to functional categories relating to host identification, detoxification and chaperone activity discussed earlier also demonstrated developmental regulation in our study. Two OR transcripts showed higher expression levels in adults than in nymphs indicating a possible sub-functionalization among the OR transcripts. Of the six chemosensory proteins (CSP) transcripts detected, two were up-regulated in adults w.r.t. nymphs. Seven cytochrome P450 transcripts including CYP6CY3 (whose over-expression increased nicotine resistance in *M. persicae*) (65) showed higher or specific expression in adults. Developmental regulation of cytochrome P450s has been shown earlier in *M. persicae* as well (27). Two GST and five SOD transcripts were found to be specific to adults. Details of these transcripts are provided in S4Table. Expression of HSPs is also known to be developmentally regulated in insects (73, 74). Twenty-one Hsp transcripts showed higher/specific expression in adult aphids in this study (S3 Table). These observations indicate that aphid adults and nymphs might employ different strategies to interact with their hosts, which is also an important factor for aphid control programmes.

#### Developmental regulation of putative effectors in L. erysimi

Some of the putative effectors discussed earlier (S5Table) were also found to be developmentally regulated. Sixteen putative effector transcripts were up-regulated in nymphs, eleven were up-regulated in adults and 34 transcripts were specific to adults (Table 5). These observations are in contrast to an earlier study by Elzinga et al (75), wherein no significant differences in expression of *M. persicae* effectors viz, *MpC002*, *Mp1*, *Mp55*, *Mp56*, *Mp57* and *Mp58*was found between adults and nymphs. Though developmental regulation of effectors has not been reported in aphids, however they are known to be developmentally regulated in other organisms viz., Gall midge (76) and nematodes (77). Considering the differential regulation seen in other feeding-related transcripts, it is possible that aphid effectors might also be specific to the developmental stage. Differential expression of putative effectors between nymphs and adults is being reported for the first time in our study.

**Table 5:**
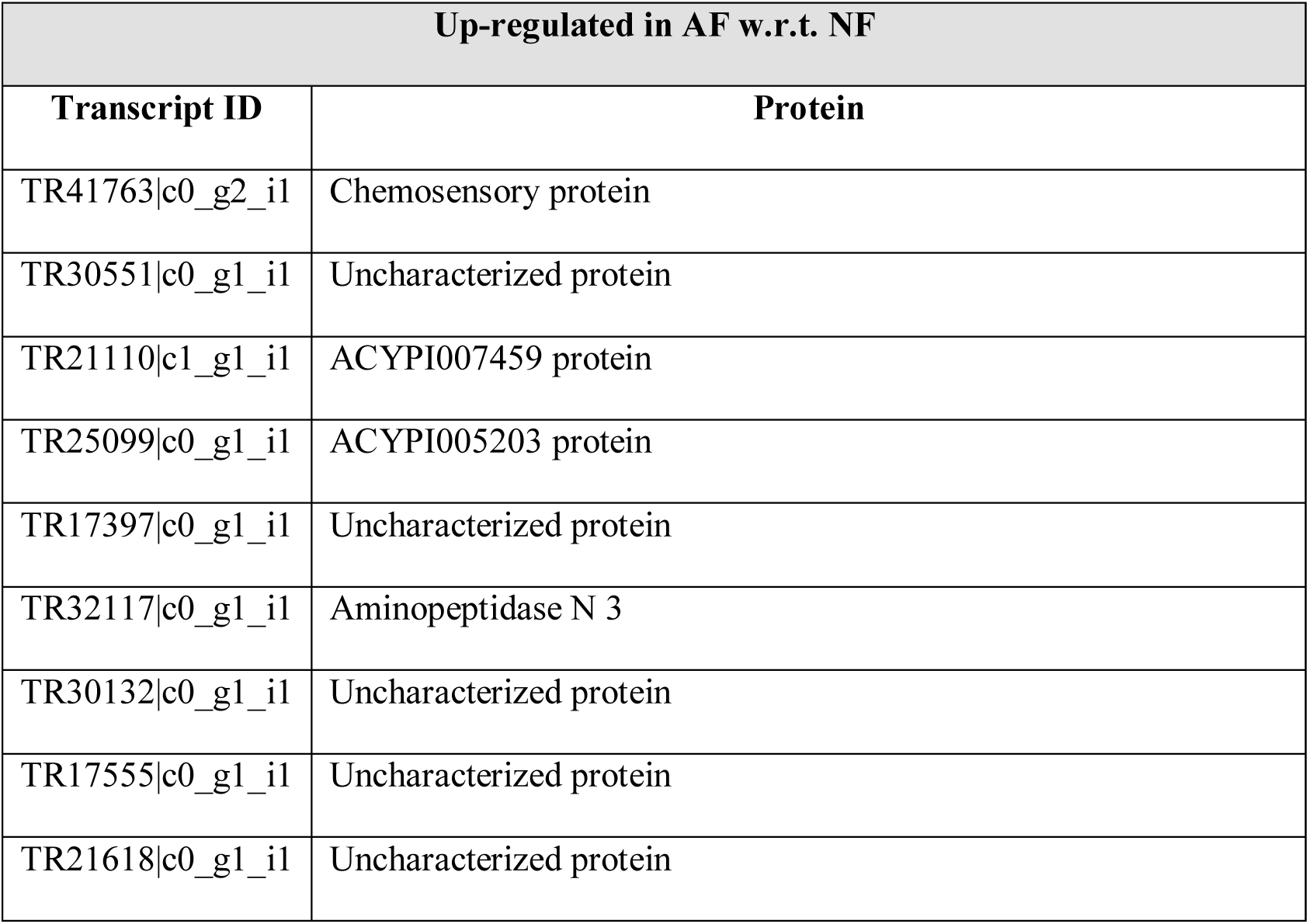
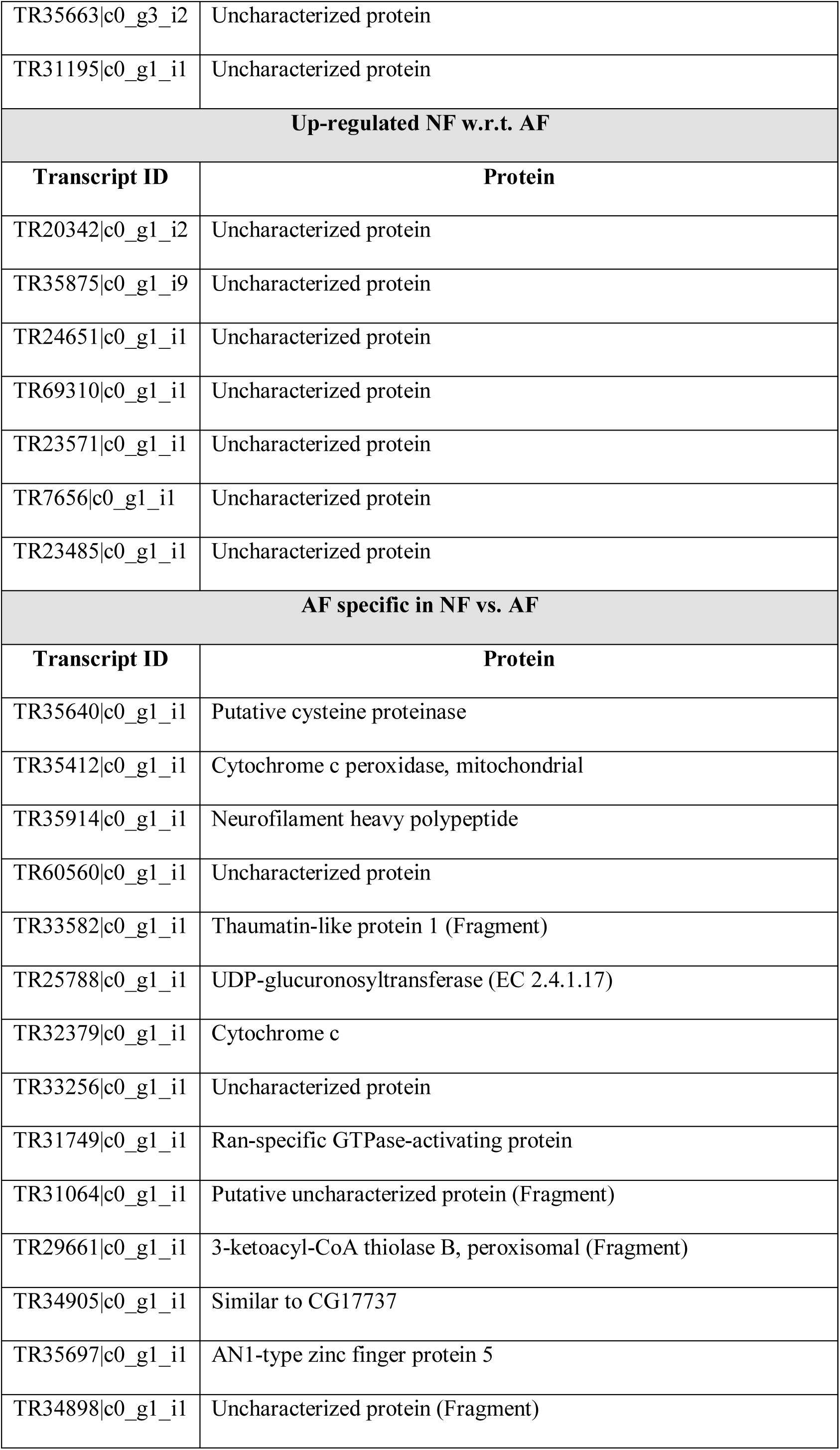
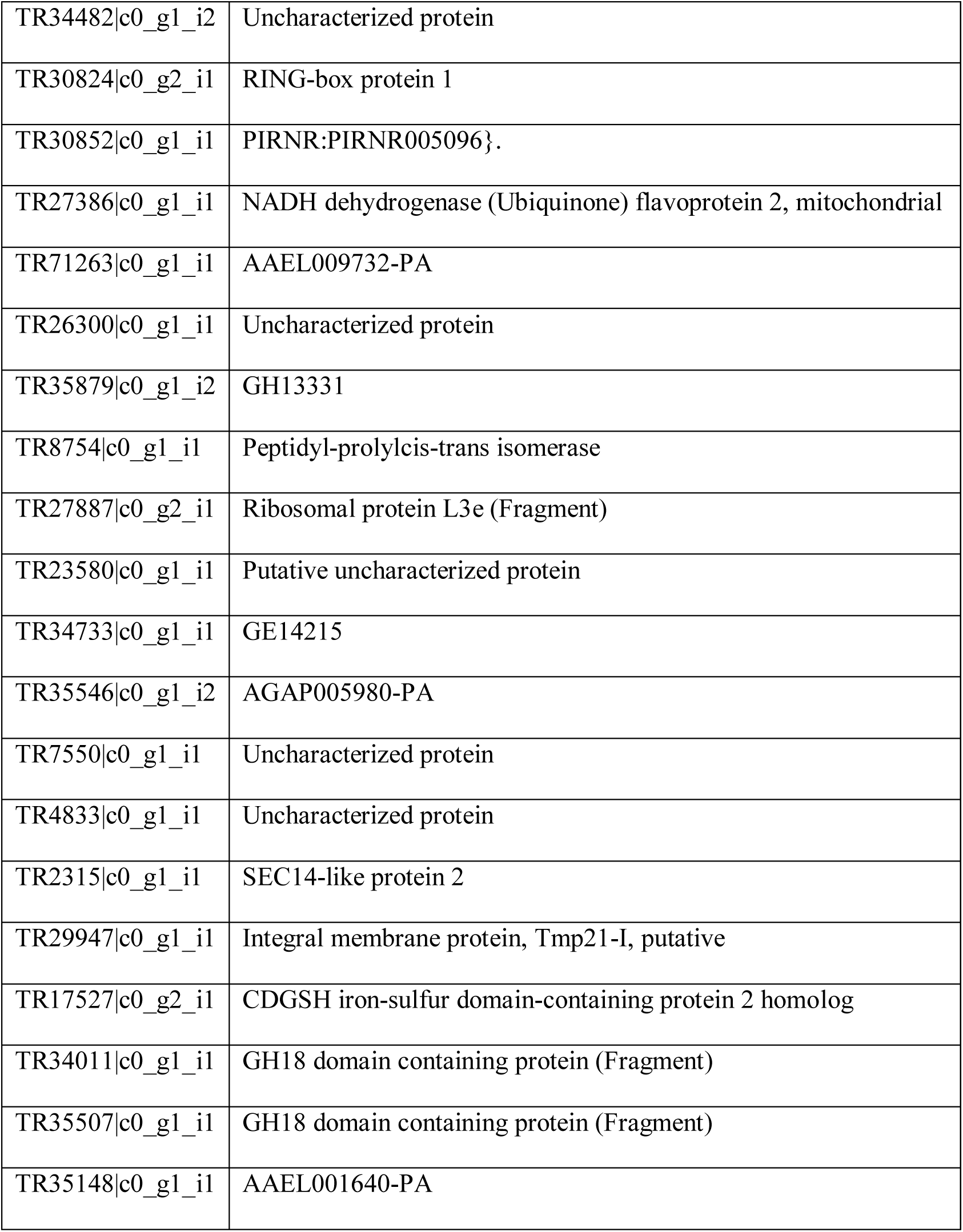
Developmentally regulated putative effectors of *L. erysimi*.

#### Differential regulation of putative developmental genes in L. erysimi

Aphid developmental genes have been extensively studied in *A. pisum.* Shigenobu et al. (78) described the components of developmental processes and signalling pathways. Brisson et al. identified major wing development genes of *Drosophila* in *A. pisum*. We analysed the occurrence and differential expression of these genes in *L. erysimi*. Out of seven major signalling pathways (Wnt, Notch, Transforming Growth Factor beta (TGFβ), Hedgehog (Hh), Receptor Tyrosine Kinase (RTK), Janus kinase/Signal Transducer, Activator of Transcription (JAK/STAT) and nuclear receptor) that were identified earlier (78) in pea aphid, we identified at least one component for each pathway in *L. erysimi*, except for the nuclear receptor pathway. The detected components of these signalling pathways and their expression profiles are provided in S6Table. Eleven transcripts of these signaling pathway components showed differential expression between nymphs and adults. Apart from the reported Notch pathway components, probable regulators such as transmembrane protein Presenilin (79), Notchless (80), the downstream regulator Protein strawberry notch (81) and the positive regulator, Enhancer of rudimentary homologue (82), were also identified. Genes involved in different developmental processes in pea aphid (78) and also detected in the *L. erysimi* transcriptome are given in S7Table. Among the wing development genes identified by Brissonet al. (30) in *A. pisum*, *engrailed*, *hedgehog*, *cubitus interruptus* and *ultrabithorax* were detected in *L. erysimi*. Expression of other development-related proteins such as, dumpy isoforms O and Z, Shaggy isoform P, strawberry notch, notch-like and DSCAM splice variants 4.1 and 4.3 was significantly higher in nymphs in comparison to adults. Shaggy is a Glycogen synthase kinase-3β (GSK-3β) homolog in *D. melanogaster* that is important for segment polarity during embryogenesis (83). It is involved in Wnt signalling (78) and follows circadian rhythm in *A. pisum* (84). DSCAM is involved in development of neuronal circuit and its orthologs have been studied in *A. pisum* as well (85). Transcripts of 11 known developmental genes from *D. melanogastor*, which have not yet been reported in aphids till now, were detected in the *L. erysimi* transcriptome. Six of these transcripts showed differential expression among the studied samples (S8Table). To the best of our knowledge, this is the first study to report differential expression of developmental genes at different stages of *L. erysimi* and correlate the expression profiles with different feeding conditions. These genes can also be potential RNAi targets.

Among the 23,532 total unannotated transcripts obtained, 20,928 transcripts were differentially regulated in AF vs. ANF, whereas 20,079 transcripts were differentially regulated in NF vs. AF (S12 Data sheet). These might represent novel transcripts in aphids or transcripts specific to *L. erysimi* and need to be analyzed for their functional roles.

### Validation of differentially expressed transcripts by qRT-PCR

For experimental validation of the digital expression data, we selected twelve transcripts showing significant differential regulation (i.e., with Log_2_ Fold change ≥2 and ≤-2) across different samples. Details of the selected transcripts and their expression pattern that were validated by qPCR have been shown in table 6. The primer details of these transcripts are given in S1 Table. The pea aphid *Elongation Factor 1α* (*EF1α*), which showed stable expression levels in all the three samples of the current study, was used as an endogenous control for normalization. Total RNA of the three samples under study was isolated in three biological replicates.

**Table 6:** Details of expression patterns of transcripts validated by qPCR.

| Transcript ID | Protein annotation | Samples compared | Digital expression pattern | qPCR expression pattern |
| --- | --- | --- | --- | --- |
| TR35875\|c0_g1_i5 | Dumpy, isoform Z | NF vs AF | Upregulated | Upregulated |
| TR35875\|c0_g1_i8 | Dumpy, isoform O | NF vs AF | Upregulated | Upregulated |
| TR35875\|c0_g1_i1 | Dumpy, isoform Z | NF vs AF | Upregulated | Upregulated |
| TR35875\|c0_g1_i11 | Putative uncharacterized protein-NFvsAF | NF vs AF | Upregulated | Upregulated |
| TR27570\|c0_g2_i1 | Lipase | NF vs AF | Upregulated | Upregulated |
| TR20342\|c0_g1_i2 | Uncharac protein | NF vs AF | Upregulated | Upregulated |
| TR35875\|c0_g1_i8 | Dumpy, isoform O | AF vs ANF | Downregulated | Downregulated |
| TR60068\|c0_g1_i1 | Projectin short variant (Fragment) | AF vs ANF | Downregulated | Downregulated |
| TR41759\|c0_g1_i1 | TCP-1/cpn60 chaperonin family protein | AF vs ANF | Downregulated | Downregulated |
| TR34264\|c0_g1_i3 | Neural-cadherin (Cadherin-N) (dN-cadherin) | AF vs ANF | Downregulated | Downregulated |
| TR3707\|c0_g1_i1 | Putative aquaporin aqpan.g (Fragment) | AF vs ANF | only in AF | Upregulated |
| TR44366\|c0_g1_i1 | Mothers against decapentaplegic homolog | NF vs AF | only in NF | Upregulated |
| TR36031\|c1_g1_i1 | Putative beta-spectrin (Fragment) | NF vs AF | only in NF | Upregulated |

Pairwise comparisons of expression profiles of these randomly selected transcripts were performed by qPCR as shown in table 6. The acquired results have been depicted in figure 5. The results of qRT-PCR analysis for each selected target were in consonance with the digital expression data (Fig5, Table 6). Three transcripts that were observed to be specific to AF (w.r.t. ANF) and NF (w.r.t. AF), viz putative aquaporin, MAD homolog and Putative beta-spectrin respectively, were seen to be upregulated in qPCR. As explained earlier, transcripts observed to be digitally specific to a sample might be simply up-regulated biologically. A putative beta-spectrin, Lipase and developmentally important transcripts, Dumpy isoform O, both variants of isoform Z and MAD homolog were up-regulated in nymphal stage w.r.t. adult stage. Dumpy isoform O, Projectin short variant, neural cadherin, TCP-1 and chaperonin showed higher expression in Adult Non-Feeding w.r.t. Adult Feeding samples thereby validating starvation stress-induced up-regulation of developmental genes, structural protein and chaperone. The potential roles of these transcripts need to be further validated experimentally.

**Fig 5.**
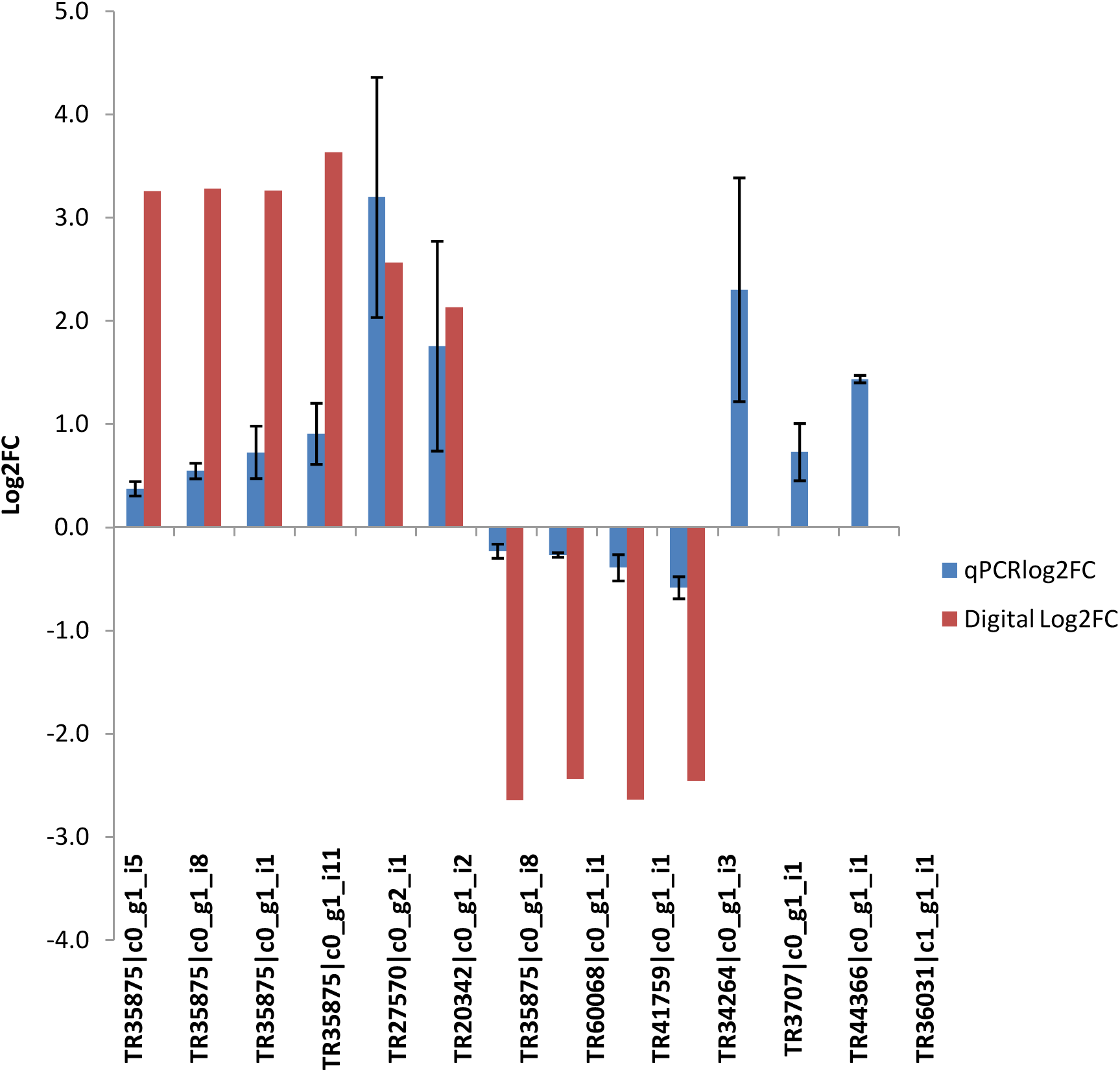
Validation of the transcriptome data using qPCR. Log_2_ fold changes of selected differentially expressed transcripts by qPCR. Levels of expression of the selected transcripts were quantified by qPCR. Relative fold changes were calculated using ΔΔCt method, comparing Adult Feeding (AF), Nymph Feeding (NF) and Adult Non-Feeding (ANF) pairwise. The mean of three independent biological replicates and their respective standard errors are represented. Label on x-axis represent the transcript IDs.

## Concluding remarks

The transcriptome data generated in this study augments existing genomic resources and identified for the first time, variations in global expression patterns of genes during feeding and non-feeding conditions in any aphid. Several of these differentially expressed transcripts are involved in host recognition, detoxification, defence response, nutrition and development indicating significant differences and similarities in aphid physiology and behaviour under the studied conditions. Several transcripts with established developmental roles in other insects are being reported for the first time in aphids in this study. Variations in expression patterns of different paralogs of genes have also been identified. These differentially regulated transcripts that are important for feeding and development can be utilized for designing effective control strategies for *L. erysimi*. Many of the unannotated transcripts, which showed significant differential regulation during feeding or development, need to be functionally characterized to improve our understanding of aphid biology. Keeping in view the lack of technical replicates in sequencing the findings of this study requires further investigation. Despite this shortcoming, it provides valuable information as the first transcriptome-based study of *L. erysimi*, a non-model aphid and a devastating pest of oilseed Brassicas in the Indian subcontinent. The resources generated in this study would facilitate basic and applied research on aphid biology, evolution and control.

## Supporting information

S1 Data sheet

S2 Data sheet

S3 Data sheet

S4 Data sheet

S5 Data sheet

S6 Data sheet

S7 Data sheet

S8 Data sheet

S9 Data sheet

S10 Data sheet

S11 Data sheet

S12 Data sheet

## ACKNOWLEDGEMENT

We thank Prof. A. K. Singh, Department of Zoology, University of Delhi, for providing us apterous adults of *L. erysimi* to initiate our population. RC is thankful for research fellowships from the University Grants Commission and Department of Biotechnology, Government of India.

## AUTHOR CONTRIBUTIONS

AJ and MA conceived the research program and supervised the research work. AJ, MA and RC designed the experiment. RC did the aphid rearing, sample collection, RNA isolation, preparation of transcriptome libraries, qPCR validation of transcripts and downstream data analyses. KK and MV did the annotation, differential expression study and signal peptide and protein localization predictions. GJ performed quality check of raw sequences. RC and AJ wrote the manuscript. MA, SG and AK provided critical inputs during experiments and preparation of manuscript.

## Supporting information

**S1 Table.** Details of primers used for qPCR validation.

**S2 Table.** Top 10 species distribution of the BLASTx against SwissProt insect sequences.

**S3 Table.** Transcript hits with molecular chaperones.

**S4 Table.** Transcripts involved in host selection, detoxification and defence and their differential expression.

**S5 Table.** Differentially regulated transcripts of putative effectors in *L. erysimi* with annotation.

**S6 Table.** Transcripts involved in Developmental signaling pathways.

**S7 Table.** Transcripts involved in Developmental processes.

**S8 Table.** Developmental genes of *L. erysimi* not reported in other aphids.

**S1 Data sheet.** Transcripts annotated by BLASTx against SwissProt insect sequences.

**S2 Data sheet.** Endosymbiont sequences from Endosymbiont BLASTn and All-organism nr-DataBase BLASTx.

**S3 Data sheet.** Transcripts annotated by BLASTx against all-organism nr DataBase.

**S4 Data sheet.** Up-regulated in Adult Feeding vs. Adult Non-Feeding.

**S5 Data sheet.** Down-regulated in Adult Feeding vs. Adult Non-Feeding.

**S6 Data sheet.** Adult Feeding specific in Adult Feeding vs. Adult Non-Feeding.

**S7 Data sheet.** Adult Non-Feeding specific in Adult Feeding vs. Adult Non-Feeding.

**S8 Data sheet.** Up-regulated in Nymph Feeding vs. Adult Feeding.

**S9 Data sheet.** Down-regulated Sequences in Nymph Feeding vs Adult Feeding.

**S10 Data sheet.** Sequences specific to Nymph-Feeding in Nymph Feeding vs Adult Feeding.

**S11 Data sheet.** Sequences specific to Adult-Feeding in Nymph Feeding vs Adult Feeding.

**S12 Data sheet.** Differential expression of unannotated transcripts.

## REFERENCES

1. Remaudière, G. & Remaudière, M. (1997). Catalogue des Aphididae du Monde. (Paris: INRA).

2. Blackman, R. L & Eastop, V. F. (2000). Aphids on the World’s Crops: An Identification and Information Guide. 2nd ed. (Chichester: Wiley).

3. Grover, A. & Pental, D. (2003). Breeding objectives and requirements for producing transgenics for major field crops of India. Current Science, 84(3).

4. Vikas, Singh, S. P., Singh, H., Hegde, D. M. & Tahir, T.A. (2007). Past progress, present scenario, nutritional value and strategies to enhance yield potential of rapeseed-mustard: an overview. Indian j Crop Science. 2(2), 245–257.

5. Bhakhetia, D. R. C. (1984). Chemical control of *Lipaphis erysimi* (Kalt.) on rapeseed and mustard crops. Punjab j. Res. Punjab Agric. Univ. 21 (63).

6. Verma, S. N. & Singh, O.P. (1987). Estimation of avoidable losses to mustard by aphid, *Lipaphis erysimi* in Madhya Pradesh. Indian j Plant Prot.15 (87).

7. Banerjee, S., Hess, D., Majumdar, P., Roy, D. & Das, S. (2004). The interactions of *Allium sativum* leaf agglutinin with a chaperonin group of unique receptor protein isolated from a bacterial endosymbiont of the mustard aphid. J. Biol. Chem.279, 23782–23789.

8. Hardie, J. (1989). Spectral specificity for targeted flight in the black bean aphid, Aphis fabae. J. of Insect Physiology. 35, 619–626.

9. Nottingham, S. F. & Hardie, J. (1993). Flight behaviour of the black bean aphid, *Aphis fabae*, and the cabbage aphid, *Brevicoryne brassicae*, in host and non-host plant odour. Physiological Entomology. 18, 389–394.

10. Park, K. C. & Hardie, J. (2004). lectrophysiological characterisation of olfactory sensilla in the black bean aphid, Aphis fabae. J. of Insect Physiology. 50, 647–655.

11. Tjallingii, W. F. (2006). Salivary secretions by aphids interacting with proteins of phloem wound responses. J. Exp. Bot. 57, 739–745.

12. Jenks, M. A., Eigenbrode, S. D. & Lemieux, B. (2002). “Cuticular Waxes of *Arabidopsis*”. The Arabidopsis Book / American Society of Plant Biologists. 1. doi: 10.1199/tab.0016.

13. Wagner, G. J., Wang, E. and Shepherd, W. (2004). New approaches for studying and exploiting an old protuberance, the plant trichome. Annal. Bot. 93, 3–11.

14. Kuśnierczyk, A., Winge, P., Jørstad, T.S., TroczyTroczyńska, J., Rossiter, J. T. and Bones, A. M. (2008). Towards global understanding of plant defence against aphids-timing and dynamics of early Arabidopsis defence responses to cabbage aphid (*Brevicoryne brassicae*) attack. Plant Cell Environ.31, 1097–1115.

15. Bones, A. M. & Rossiter, J. T. (1995). “Glucosinolates in cruciferous crops”, New horizons in oilseed rape, ed. D. H. Scarisbrick and A. J. Ferguson (Cambridge: Semundo), 46–67.

16. Harrison, R.L. & Bonning, B.C. (2010). Proteases as Insecticidal Agents. Toxins. 2(5), 935–953.

17. Wieczorek, M., Otlewski, J., Cook, J., Parks, K., Leluk, J. (1985). The squash family of serine proteinase inhibitors. Amino acid sequences and association equilibrium constants of inhibitors from squash, summer squash, zucchini, and cucumber seeds. Biochem. Biophys. Res. Commun.126, 646–652.

18. Gaupels, F., Knauer, T. and van Bel, A.J.E. (2008). A combinatory approach for analysis of protein sets in barley sieve-tube samples using EDTA facilitated exudation and aphid stylectomy. J. Plant Physiol.165, 95–103.

19. Hao, P., Liu, C., Wang, Y., Chen, R., Tang, M., Du, B., et al. (2008). Herbivore-induced callose deposition on the sieve plates of rice: an important mechanism for host resistance. Plant Physiol.146,1810–1820.

20. da Silva, S. E. B., Franca, J. F. & Pareja, M. (2016). Olfactory response of four aphidophagous insects to aphid- and caterpillar-induced plant volatiles. Arthropod-Plant Interactions.10, 331–340.

21. Tjallingii, W.F. and Esch, T.H. (1993).Fine structure of aphid stylet routes in plant tissues in correlation with EPG signals. Physiol. Entomol.18, 313–328.

22. Miles, P.W. (1990). Aphid salivary secretions and their involvement in plant toxicoses in Aphid-plant genotype interactions, ed. R. K. Campbell & R. D. Eikenbary (Amsterdam: Elsevier),131–147.

23. Navazio, L., Moscatiello, R., Bellincampi, D., Baldan, B., Meggio, F., Brini, M., et al. (2002). The role of calcium in oligogalacturonide-activated signalling in soybean cells. Planta. 215, 596–605.

24. Will, T., Tjallingii, W. F., Thonnessen, A. & van Bel, A. J. E. (2007). Molecular sabotage of plant defence by aphid saliva. Proc. Natl. Acad. Sci. USA.104, 10536–10541.

25. Carolan, J. C., Caragea, D., Reardon, K.T., Mutti, N. S., Dittmer, N., Pappan, K., et al. (2011). Predicted effector molecules in the salivary secretome of the pea aphid (*Acyrthosiphonpisum*): a dual transcriptomic/proteomic approach. J. Proteome Res.10, 1505–1518.

26. Bridges, M., Jones, A. M. E., Bones, A. M., Hodgson, C., Cole, R., Bartlet, E., et al. (2002). Spatial organization of the glucosinolate–myrosinase system in brassica specialist aphids is similar to that of the host plant. Proc. R. Soc. Lond. B. 269, 187–191.

27. Ji, R., Wang, Y., Cheng, Y., Zhang, M., Zhang, H-B., Zhu, L., Fang, J. and Zhu-Salzman, K. (2016). Transcriptome analysis of green peach aphid (*Myzus persicae*): Insight into developmental regulation and inter-species divergence. Front. Plant Sci. 7: 1562.doi: 10.3389/fpls.2016.01562.

28. Muller, C. B., Williams, I. S. & Hardie, J. (2001). The role of nutrition, crowding and interspecific interactions in the development of winged aphids. Ecological Entomology.26, 330–340.

29. Kati, A. and Hardie, J. (2010). Regulation of wing formation and adult development in an aphid host, Aphis fabae, by the parasitoid Aphidius colemani. J. Insect Physiology.56, 14–20.

30. Brisson, J. A., Ishlkawa, A. & Mlura, T. (2010).Wing development genes of the pea aphid and differential gene expression between winged and unwinged morphs. Insect Molecular Biology, 19, 63–73.

31. Legeai F, Rizk G, Walsh T, Edwards O, Gordon K, Lavenier D, et al. (2010). Bioinformatic prediction, deep sequencing of microRNAs and expression analysis during phenotypic plasticity in the pea aphid, *Acyrthosiphon pisum*. BMC Genomics. 11, 281.

32. Duncan, E. J., Leask, M. P. and Dearden, P. K. (2013). The pea aphid (*Acyrthosiphon pisum*) genome encodes two divergent early developmental programs. Developmental Biology. 377, 262–274.

33. Hansen, A. K. and Moran, N. A. (2011). Aphid genome expression reveals host–symbiont cooperation in the production of amino acids. Proc. Natl. Acad. Sci. USA. 108, 2849–2854.

34. Liu, S., Chougule, N. P., Vijayendran, D., Bonning, B. C. (2012). Deep sequencing of the transcriptomes of soybean aphid and associated endosymbionts. PLoSONE. 7(9), e45161; 10.1371/journal.pone.0045161.

35. Wang, D., Li, Q., Jones, H. D., Bruce, T. & Xia, L. (2014). Comparative transcriptomic analyses revealed divergences of two agriculturally important aphid species. BMC Genomics. 15, 1023.

36. Li ZQ, Zhang S, Luo J-Y, Wang C-Y, Lv L-M, Dong S-L, et al.(2013). Ecological adaption analysis of the cotton aphid (*Aphis gossypii*) in different phenotypes by transcriptome comparison. PLoSONE.8(12), e83180; 10.1371/journal.pone.0083180.

37. Coppola, V., Coppola, M., Rocco, M., Digilio, M.C., D’Ambrosio, C., Renzone, G., et al. (2013). Transcriptomic and proteomic analysis of a compatible tomato-aphid interaction reveals a predominant salicylic acid-dependent plant response. BMC Genomics. 14, 515, doi: 10.1186/1471-2164-14-515.

38. Dubey, N. K., Goel, R., Ranjan, A., Idris, A., Singh, S. K., Bag, S. K., et al. (2013). Comparative transcriptome analysis of *Gossypium hirsutum* L. in response to sap sucking insects: aphid and whitefly. BMC Genomics. 14, 241.

39. Jaouannet, M., Morris, J. A., Hedley, P. E., Bos, J. I. B. (2015). Characterization of *Arabidopsis* transcriptional responses to different aphid species reveals genes that contribute to host susceptibility and non-host resistance. PLoS Pathog. 11:5. doi: 10.1371/journal.ppat.1004918.

40. Chomczynski, P. & Sacchi, N. (1987). Single-step method of RNA isolation by acid guanidiumthiocyanate-phenol-chloroform extraction. Analytical Biochemistry.162, 156–159.

41. Patel, R. K. & Jain, M. (2012). NGS QC Toolkit: A Toolkit for Quality Control of Next Generation Sequencing Data. PLoS ONE. 7(2), e30619; 10.1371/journal.pone.0030619.

42. Haas, B. J., Papanicolaou, A., Yassour, M., Grabherr, M., Blood, P. D., Bowden, J., et al. (2013). *De novo* transcript sequence reconstruction from RNA-Seq: reference generation and analysis with Trinity. Nature Protocols. 8, 1494–1512.

43. Langmead, B. and Salzberg, S. L. (2012). Fast gapped-read alignment with Bowtie 2. Nature Methods.9, 357–359.

44. Li, B. and Dewey, C. N. (2011). RSEM: accurate transcript quantification from RNA-Seq data with or without a reference genome. BMC Bioinformatics. 12, 323.

45. Bankar, K. G., Todur, V. N., Shukla, R. N. and Vasudevan, M. (2015). Ameliorated de novo transcriptome assembly using Illumina paired end sequence data with Trinity Assembler. Genomics Data. 5, 352–359.

46. Li, W. and Godzik, A. (2006). Cd-hit: a fast program for clustering and comparing large sets of protein or nucleotide sequences. Bioinformatics.1, 1658–1659.

47. Anders, S. & Huber, W. (2010). Differential expression analysis for sequence count data. Genome Biology. 11, R106.

48. Altschul, S. F., Gish, W., Miller, W., Myers, E. W. and Lipman, D. J. (1990). Basic local alignment search tool. J. Mol. Biol. 215(3), 403–410.

49. Livak, K. J. & Schmittgen, T. D. (2001). Analysis of relative gene expression data using real-time quantitative PCR and the 2(-Delta Delta C(T) Method. Methods. 25(4), 402–408.

50. Tjallingii, W. F. (1990). Continuous recording of stylet penetration activities by aphids, in Aphid-Plant Genotype Interactions, ed. R. K. Campbell & R. D. Eikenbary (Amsterdam: Elsevier Science Publishers B.V.), 89–99.

51. van Emden, H.F.(1977). Failure of the aphid, *Myzus persicae*, to compensate for poor diet during early growth. Physiological Entomology. 2, 53–58.

52. . The International Aphid Genomics Consortium Genome. (2010). Sequence of the Pea Aphid Acyrthosiphon pisum. PLoS Biol. 8: 2.doi: 10.1371/journal.pbio.1000313.

53. Nicholson S J, Nickerson ML, Dean M, Song Y, Hoyt PR, Rhee H, et al.(2015). The genome of *Diuraphis noxia*, a global aphid pest of small grains. BMC Genomics.16, 429, doi: 10.1186/s12864-015-1525-1.

54. Will, T., Schmidtberg, H., Skaljac, M. and Vilcinskas, A. (2017). Heat shock protein 83 plays pleiotropic roles in embryogenesis, longevity and fecundity of the pea aphid *Acyrthosiphon pisum*. Dev. Genes Evol. 227, 1–9.

55. Dunbar, H. E., Wilson, A. C. C., Ferguson, N. R. & Moran, N. A. (2007). Aphid thermal tolerance is governed by a point mutation in bacterial symbionts. PLoS Biol.5:5. doi: 10.1371/journal.pbio.0050096.

56. Ma, R., Reese, J. C., Black IV, W. C. and Bramel-Cox, P. (1990). Detection of pectinesterase and polygalacturonase from salivary secretions of living greenbugs, *Schizaphis graminum* (Homoptera: Aphididae). J. Insect Physiol. 36, 507–512.

57. Cai, Q-N., Han, Y., Cao, Y-Z., Hu, Y., Zhao, X. and Bi, J-L. (2009). Detoxification of gramine by the cereal aphid *Sitobion avenae*. J. Chem. Ecol. 35, 320–325.

58. Lei, J. and Zhu-Salzman, K. (2015). Enhanced aphid detoxification when confronted by a host with elevated ROS production. Plant signalling & behaviour. 10:4. doi: 10.1080/15592324.2015.1010936.

59. Martins, D., Kathiresan, M. & English, A. M. (2013). Cytochrome c peroxidase is a mitochondrial heme-based H2O2 sensor that modulates antioxidant defence. Free Radical Biology and Medicine. 65, 541–551.

60. Deng, F. & Zhao, Z. (2014). Influence of catalase gene silencing on the survivability of Sitobion avenae. Archives of insect biochemistry and physiology. 86(1), 46–57.

61. Sekelsky, J. J., Newfeld, S. J., Raftery, L. A., Chartoff, E. H. & Gelbart, W. M. (1995). Genetic characterization and cloning of *Mothers against dpp*, a gene required for *decapentaplegic* function in *Drosophila melanogaster*. Genetics. 139, 1347–1358.

62. Lamb, R. J. and MacKay, P. A. (1983). “Micro-evolution of the migratory tendency, photoperiodic response and developmental threshold of the pea aphid, *Acyrthosiphon pisum*,” in Diapause and life-cycle strategies in insects, ed. V. K. Brown and I. Hodek (The Hague: Dr. W. Junk Publishers), 210–217.

63. Weisser, W. W. and Braendle, C. (2001). Body colour and genetic variation in winged morph production in the pea aphid. Entomologia Experimentalis et Applicata. 99, 217–223.

64. Pelosi, P., Zhou, J-J., Ban, L.P. and Calvello, M. (2006). Soluble proteins in insect chemical communication. Cell. Mol. Life Sci. 63, 1658–1676.

65. Bass, C., Puinean, A. M., Zimmer, C. T., Denholm, I., Field, L. M., Foster, L. M., et al. (2014). The evolution of insecticide resistance in the peach potato aphid, *Myzus persicae*. Insect Biochemistry and Molecular Biology. 51, 41–51, doi: 10.1016/j.ibmb.2014.05.003.

66. Prasain, K., Nguyen, T. D. H., Gorman, M. J., Barrigan, L. M., Peng, Z., Kanost, M. R., et al. (2012). Redox potentials, laccase oxidation, and antilarval activities of substituted phenols. Bioorg. Med. Chem.20(5), 1679–1689.doi: 10.1016/j.bmc.2012.01.021.

67. Bansal, R., Mian, M. A. R., Mittapalli, O. & Michel, A. P. (2014). RNA-Seq reveals a xenobiotic stress response in the soybean aphid, *Aphis glycines*, when fed aphid-resistant soybean. BMC Genomics.15, 972.

68. Nakabachi, A., Shigenobu, S. and Miyagishima, S. (2010). Chitinase-like proteins encoded in the genome of the pea aphid, *Acyrthosiphon pisum*. Insect Molecular Biology. 19 (Suppl. 2), 175–185.

69. Nicholson, S. J., Hartson, S. D. and Puterka, G. J. (2012). Proteomic analysis of secreted saliva from Russian wheat aphid (*Diuraphis noxia* Kurd.) biotypes that differ in virulence to wheat. J Proteomics. 75(7), 2252–68. doi: 10.1016/j.jprot.2012.01.031.

70. Rao, S. A. K., Carolan, J. C. & Wilkinson, T. L.(2013). Proteomic profiling of cereal aphid saliva reveals both ubiquitous and adaptive secreted proteins. PLoS ONE.8: 2. doi: 10.1371/journal.pone.0057413.

71. Orii, K. E., Lee, Y., Kondo, N. & McKinnon, P. J.(2006).Selective utilization of nonhomologous end-joining and homologous recombination DNA repair pathways during nervous system development. Proc. Natl. Acad. Sci. USA. 103, 10017–10022.

72. Douglas, A. E. and van Emden, H. F. (2007). Nutrition and symbiosis, in Aphids as crop pests, ed. H. F. van Emden and R. Harrington (Cabi), 115–134.

73. Tachibana, S-I, Numata, H. and Goto, S. G. (2005). Gene expression of heat-shock proteins (Hsp23, Hsp70 and Hsp90) during and after larval diapause in the blow fly Lucilia sericata. J. of Insect Physiology. 51, 641–647.

74. Mahroof, R., Zhu, K. Y., Neven, L., Subramanyam, B. and Bai, J. (2005). Expression patterns of three heat shock protein 70 genes among developmental stages of the red flour beetle, *Tribolium castaneum* (Coleoptera: Tenebrionidae). Comparative Biochemistry and Physiology. Part A. 141, 247 – 256.

75. Elzinga, D. A., De Vos, M. and Jander, G. (2014). Suppression of plant defences by a *Myzus persicae* (green peach aphid) salivary effector protein. Mol. Plant Microbe Interact. 27(7), 747–756.

76. Chen, M-S., Liu, S., Wang, H., Cheng, X., Bouhssini, M. E. and Whitworth, R. J. Massive shift in gene expression during transitions between developmental stages of the gall midge, *Mayetiola destructor*. PLoS ONE. 11:5. doi: 10.1371/journal.pone.0155616.

77. Mitchum, M. G. et al. (2013). Nematode effector proteins: an emerging paradigm of parasitism. New Phytologist. 199, 879–894. doi: 10.1111/nph.12323.

78. Shigenobu, S., Bickel, R. D., Brisson, J. A., Butts, T., Chang, C. C., Christiaens, et al. (2010). Comprehensive survey of developmental genes in the pea aphid, *Acyrthosiphon pisum*: frequent lineage-specific duplications and losses of developmental genes. Insect Molecular Biology. 19, 47–62. doi: 10.1111/j.1365-2583.2009.00944.x

79. Struhl, G. and Greenwald, I. (1999). Presenilin is required for activity and nuclear access of Notch in *Drosophila*. Nature. 398(6727), 522–525.

80. Royet, J., Bouwmeester, T. and Cohen, S. M. (1998). Notchless encodes a novel WD40-repeat-containing protein that modulates Notch signaling activity. EMBO J.17(24), 7351–7360.

81. Majumdar, A., Nagaraj, R. and Banerjee, U. (1997). *strawberry notch* encodes a conserved nuclear protein that functions downstream of *Notch* and regulates gene expression along the developing wing margin of *Drosophila*. Genes & Development.11, 1341–1353.

82. Tsubota, S. I., Vogel, A. C., Phillips, A. C., Ibach, S. M., Weber, N. K., Kostrzebski, M. A., et al. (2011). Interactions between *enhancer of rudimentary* and *Notch* and *deltex* reveal a regulatory function of *enhancer of rudimentary* in the Notch signalling pathway in *Drosophila melanogaster*. Fly. 5:4, 275–284. doi: 10.4161/fly.5.4.17807.

83. Fiol, C. J., Williams, J. S., Chou, C. H., Wang, Q. M., Roach, P. J. & Andrisani, O. M. (1994). A secondary phosphorylation of CREB341 at Serf2’ is required for the CAMP-mediated control of gene expression: a role for glycogen synthase kinase-3 in the control of gene expression. J. Biol. Chem. 269 (51), 32187–32193.

84. Cortes, T., Ortiz-Rivas, B. and Martínez-Torres, D. (2010). Identification and characterization of circadian clock genes in the pea aphid *Acyrthosiphon pisum*. Insect Molecular Biology.19, 123–139.

85. Armitage, S. A. O., Freiburg, R. Y., Kurtz, J. and Bravo, I. G. (2012). The evolution of Dscam genes across the arthropods. BMC Evolutionary Biology. 12, 53.

